# The CaMKII D135N mutation blocks kinase activity and reduces GluN2B binding

**DOI:** 10.1101/2025.09.20.677536

**Authors:** Matthew E. Larsen, C. Madison Barker, Raul Satoshi Vargas, Steven J. Coultrap, K. Ulrich Bayer

## Abstract

Three recent studies claimed that induction of long-term potentiation (LTP) of synaptic strength requires structural rather than enzymatic functions of the Ca^2+^/Calmodulin(CaM)-dependent protein kinase II (CaMKII). One study utilized the CaMKII D135N mutation, which was claimed to abolish enzymatic activity without affecting the structural function, i.e. binding to GluN2B. We found here that the D135N mutant indeed abolished enzymatic kinase activity and autophosphorylation at T286 (pT286). However, D135N mutation additionally reduced binding to GluN2B and prevented persistence of co-condensation with GluN2B. Similar as with the T286A mutant, GluN2B binding of the D135N mutant could be partially rescued by AS283, an inhibitor that directly enhances GluN2B binding. This partial effect on GluN2B binding has to be taken into account when using D135N to distinguish between enzymatic versus structural functions of CaMKII. Nonetheless, as discussed here, the D135N mutant indeed supports a structural rather than enzymatic CaMKII function in LTP induction.

## INTRODUCTION

Learning and memory is thought to require long-term potentiation (LTP) of synaptic strength in the hippocampus, and a central dogma of molecular neuroscience that has stood undisputed for over 30 years was that LTP induction requires enzymatic activity of the Ca^2+^/calmodulin-dependent protein kinase II (CaMKII)(1–4). However, three recent studies claimed that LTP induction instead requires structural not enzymatic functions of CaMKII, specifically the regulated binding to the NMDA-type glutamate receptor subunit GluN2B (5–7). In order to arrive at this surprising conclusion, the studies used completely different sets of tools. One of the studies relied on the CaMKII D135N mutation, which was claimed to abolish enzymatic activity without affecting GluN2B binding (6). However, the results from the GluN2B binding were somewhat unclear, as the only binding assay was *in vitro* and lacked any negative control to establish baseline. One study that was cited for abolished enzymatic activity by the D135N mutation does not contain direct experimental data about enzymatic activity, but instead shows that the D135N mutant can still bind both substrate and ATP, as the study provides crystal structures of such trimeric complexes (8). Nonetheless, D135N should at least reduce enzymatic activity, as the residue is homologous to D166 in PKA, which is implicated to interact with the hydroxyl group of the substrate residue phosphorylated during the enzymatic kinase reaction (9). Indeed, D166N mutation of mammalian PKA significantly reduced kinase activity (10), and the homologous mutations in yeast PKA and mammalian PhK appear to block kinase activity almost completely, with less than 1% of residual activity reported (11, 12). Overall, the current characterization of the CaMKII D135N mutant strongly suggests at least significantly reduced CaMKII activity and likely retention of at least some GluN2B binding. However, the effect sizes remain unclear, limiting the conclusions that can be drawn from experiments with this tool mutant.

Thus, we here characterized the effects of the CaMKII D135N mutation. We found that the D135N abolished enzymatic activity and autophosphorylation at T286 (pT286). However, while the D135 mutation did not abolish GluN2B binding, it was significantly reduced. Similarly, co-condensation of CaMKII with GluN2B was also reduced, and the persistence of such co-condensates beyond removal of Ca^2+^ was eliminated. These findings need to be taken into account in the interpretation of results with this tool mutant. However, as delineated in our discussion, these findings nonetheless further support the central claim of the recent studies that reported LTP induction by structural functions of CaMKII.

## RESULTS

### The D135N mutation abolishes enzymatic CaMKII activity

In order to test for the effect of the D135N mutation on enzymatic CaMKII activity, we used an established biochemical kinase assay that measures ^32^P incorporation into a peptide substrate of CaMKII *in vitro* (13, 14). We compared activity of CaMKII wildtype, D135N, and K42M that were expressed in HEK293 cells (as GFP-fusion proteins), with extracts from mock-transfected HEK cells as additional negative control (Fig. 1A). For CaMKII wildtype, a reaction rate of ~450 reaction per min was measured, consistent with previous findings (13). The K42M mutant is kinase dead because of abolished nucleotide binding and thus completely abolished kinase activity (Fig. 1A), as expected. Importantly, activity of the D135N mutant was indistinguishable from the kinase-dead K42M mutant (Fig. 1A), indicating that the D135N mutation also abolishes kinase activity essentially completely.

**Figure 1:**
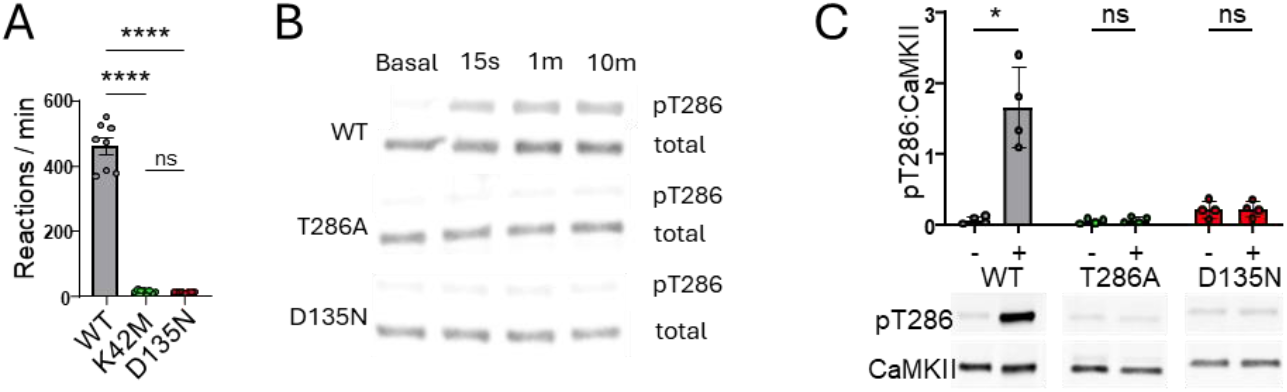
The CaMKII D135N mutation abolished enzymatic kinase activity and autophosphorylation of T286. Error bars indicate SEM. (**A**) The D135N mutation prevented Ca^2+^/CaM-induced substrate phosphorylation to the same extent as the kinase dead K42M mutation that prevents ATP binding (n=8, one-way ANOVA with Tukey’s multiple comparisons test). (**B**) Immunoblot of in vitro kinase reactions on ice in 1 mM ATP measuring purified CaMKIIα autophosphorylation at T286 after 15s, 1m, and 10m. (**C**) Representative immunoblot and quantification of in vitro kinase reactions at 30 °C with and without 1 mM ATP measuring purified CaMKIIα autophosphorylation at T286 (n=4, RM two-way ANOVA with Šídák’s multiple comparisons test). *p<0.5, ****p<0.0001

### The D135N mutation abolishes autophosphorylation of CaMKII at T286

As the D135N mutation abolishes enzymatic CaMKII activity towards an exogenous substrate (Fig. 1A), it would be expected to reduce also the CaMKII autophosphorylation at T286 (pT286). However, as this autophosphorylation is extremely fast and efficient, we decided to formally test the effect of the D135N mutation on pT286 (Fig. 1B,C). First, pT286 reactions were performed on ice, as this eliminates autophosphorylation at other residues, thereby restricting autophosphorylation to T286. Any residual detection of pT286 in the D135N mutant did not exceed the background level that was also seen before the phosphorylation reaction and in the kinase dead K42M mutant (Fig. 1B). Next, we tested pT286 reactions for 1 min at 30°C. In this case, a CaMKII T286A mutant was used as negative control. Notably, at the 1 min time point, CaMKII wildtype showed maximal pT286 even after reactions on ice (Fig. 1B). Again, the D135N mutation prevented any significant pT286 above background also when the reactions were done at 30°C (Fig. 1C).

### The CaMKII D135N mutant binds to GluN2B *in vitro*

In biochemical reactions *in vitro*, CaMKII binding to the cytoplasmic C-terminus of GluN2B (immobilized as GST-fusion protein on anti-GST coated microtiter plates) can be induced by either Ca^2+^/CaM or by pT286 (15, 16). Without pT286 or Ca^2+^/CaM, no GluN2B binding was seen in this *in vitro* assay for CaMKII wildtype or for the D135N mutant (Fig. 2). A Ca^2+^/CaM stimulus induced significant GluN2B binding for both CaMKII wildtype and for the D135N mutant (Fig. 2), similarly as described previously (6). However, in contrast to the previous report, significantly less binding was detected for the D135 mutant compared to CaMKII wildtype (Fig. 2). Notably, the binding reaction was performed in the presence of ADP, not ATP, in order to prevent two phosphorylation reactions that regulate CaMKII binding to GluN2B (16, 17): the negative-regulatory phosphorylation of GluN2B as S1303 and the positive-regulatory phosphorylation of CaMKII T286. Thus, the lower level of D135N binding cannot be explained by its lack of enzymatic activity that is required for pT286. However, despite the lower level of binding, our results demonstrate the principal capacity of the CaMKII D135N mutant to bind to GluN2B.

**Figure 2:**
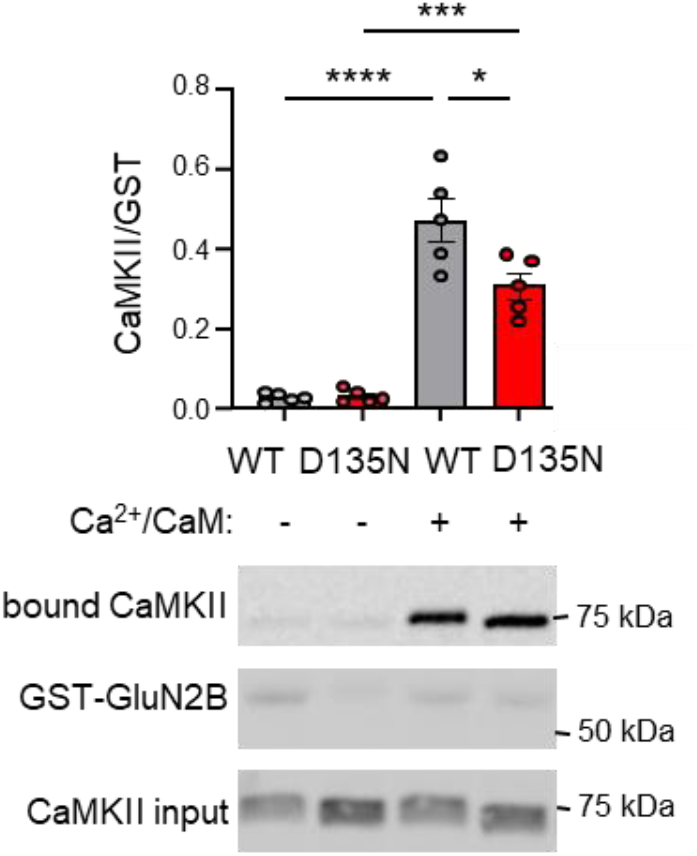
The CaMKII D135N mutation reduces binding to GluN2B *in vitro*. Representative immunoblots and quantification of in vitro binding of YFP2–CaMKII to GST– GluN2Bc immobilized on microtiter plates. Ca^2+^/CaM stimulation induced binding, with D135N showing reduced binding compared to WT–CaMKII. No binding was observed in the absence of Ca^2+^/CaM, as expected. Addition of AS283 rescued D135N binding to wild-type levels. (n=5 replicates, one-way ANOVA with Bonferroni’s post-hoc test). ns>0.5, *p<0.5

### The D135N mutation reduces GluN2B binding of paCaMKII within HEK cells

While the pT286-incompetent CaMKII T286A mutant can bind to GluN2B in biochemical assays, it shows significantly reduced GluN2B binding within HEK cells after ionomycin-induced Ca^2+^ stimuli and significantly reduced movement to excitatory synapses in hippocampal neurons in response to LTP stimuli (a movement that is mediated by GluN2B binding). Thus, as the D135N mutant is also pT286-incompetent (see Fig. 1B,C), the D135N mutation would be expected to cause similar impairments as the T286A mutation. We decided to test this first in context of the photoactivatable paCaMKII (18). Stimulation of paCaMKII with blue light induces kinase activity (18), binding to GluN2B in HEK cells (5), and movement to excitatory synapses in hippocampal neurons (5, 7). Notably, the T286A mutation abolished this paCaMKII movement in neurons, but the movement was restored by AS283 or AS397, which are ATP-competitive CaMKII inhibitors that directly enhance CaMKII binding to GluN2B and restores LTP in T286A mutant mice (5, 7). Here, we monitored movement of a GFP-labelled paCaMKII to a membrane targeted mCherry-labelled GluN2B C-tail construct in HEK cells. In order to eliminate additional regulation of the binding by GluN2B phosphorylation at S1303 (which would be induced by CaMKII wildtype and T286A, but not by the inactive D135N), a S1303A mutant GluN2B construct was used. For GFP-paCaMKII wildtype, significant light-induced co-localization with GluN2B was observed already during the initial image acquisition of a 6-image confocal z-stack (at 0.5 µm intervals) for each cell (Fig. 3A,B): The co-localization was significantly higher in the last image of each z-stack compared to the first image (Fig. 3A). This resulted in significant co-localization also in the combined projection image of these z-stacks (Fig. 3B). By contrast, no such co-localization was induced by image acquisition for the GFP-paCaMKII D135N or T286A mutants (Fig. 3B).

**Figure 3:**
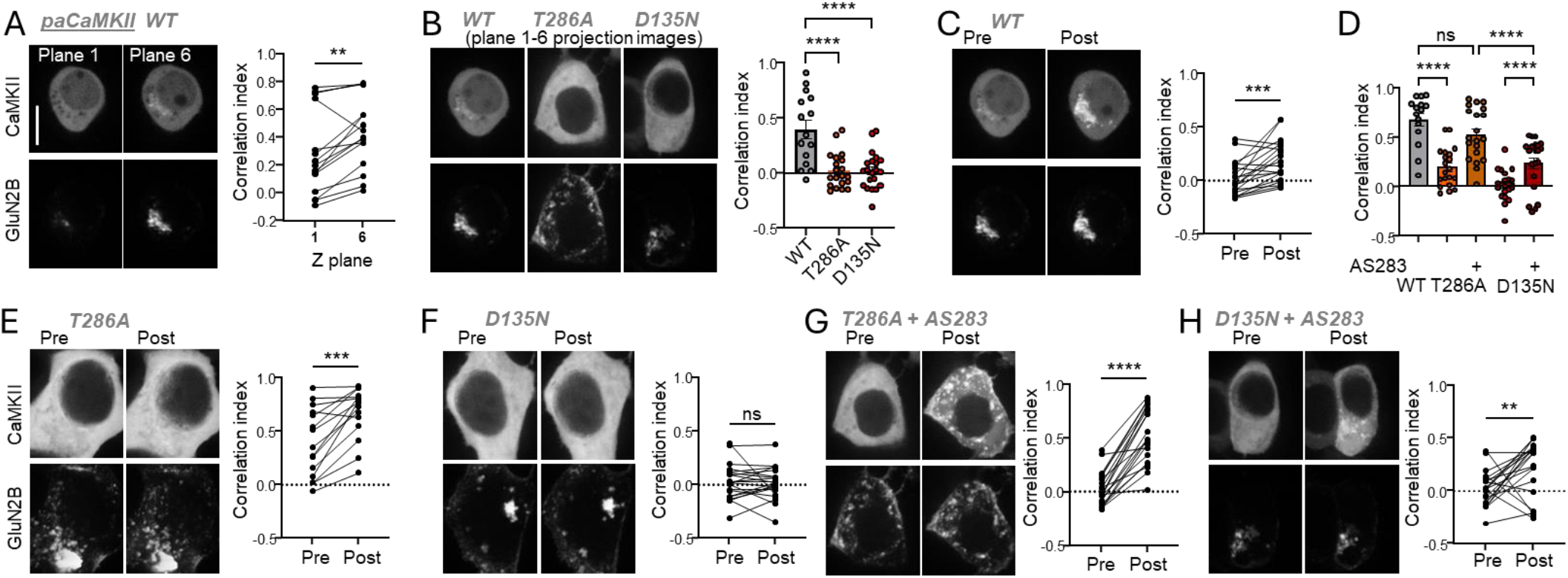
The D135N mutation reduces light-induced movement of paCaMKII to GluN2B within HEK cells. (**A**) Representative images of GFP-paCaMKII and mCh-GluN2B in HEK cells at the first and last Z-stacks of the initial time point and quantification (n=15 cells, paired t-test). (**B**) Comparison of baseline correlation for WT vs T286A and D135N (n=15,21,20 cells, one-way ANOVA with Tukey’s multiple comparisons test). (**C**) Representative images of WT GFP-paCaMKII and mCh-GluN2B pre- and post-photoactivation and quantification (n=15 cells, paired t-test). (**D**) Quantification of CaMKII-GluN2B correlation post-stimulation for WT, T286A, and D135N paCaMKII with and without AS283 (n=15,21,21,20,20 cells, two-way ANOVA with Šídák’s multiple comparison’s test). (**E**) Representative images of T286A GFP-paCaMKII and mCh-GluN2B pre- and post-photoactivation and quantification (n=20 cells, paired t-test). (**F**) Representative images of D135N GFP-paCaMKII and mCh-GluN2B pre- and post-photoactivation and quantification (n=21 cells, paired t-test). (**G**) Representative images of T286A GFP-paCaMKII and mCh-GluN2B pre- and post-photoactivation following AS283 treatment and quantification (n=20 cells, paired t-test). (**F**) Representative images of D135N GFP-paCaMKII and mCh-GluN2B pre- and post-photoactivation following AS283 treatment and quantification (n=21 cells, paired t-test). *p<0.5, **p<0.01, ***p<0.001, ****p<0.0001

Subsequent additional light stimulation resulted in a significant further increase in GluN2B co-localization for GFP-paCaMKII wildtype (Fig. 3C). A significant increase was also observed for the T286A mutant, albeit to a level that was much lower than seen for wildtype (Fig. 3D,E). By contrast, for the D135N mutant, no increase in co-localization was observed at all (Fig. 3D,F). Adding the AS283 inhibitor that enhances GluN2B binding 5 min before a second additional light stimulation caused a further increase in co-localization for the T286A, which was now similar as the co-localization normally seen with wildtype (Fig. 3D,G). For the D135N mutant, the addition of AS283 also enabled a significant increase in GluN2B co-localization in response to the second additional light stimulus; however, the co-localization still remained significantly lower compared to wildtype or T286A (Fig. 3D,H). Overall, these results demonstrate that the D135N mutation significantly reduced GluN2B binding within cells, and that this reduction can be partially but not fully rescued by AS283.

### The D135N mutation reduces Ca^2+^-induced CaMKII binding to GluN2B in HEK cells

Next we tested the effect on CaMKII movement to the same GluN2B C-tail construct in HEK cells when triggered instead by an ionomycin-induced Ca^2+^ signal (Fig. 4). In this case, the same movement seen for YFP-CaMKII WT (Fig. 4A,B) was also obtained for the GFP-tagged T286A mutant even in absence of AS283 (Fig. 4B,C) and addition of AS283 did not further increase this movement (Fig. 4B,D). By contrast, while ionomycin also induced some movement of the YFP-tagged D135N mutant, this movement was significantly reduced compared to WT (Fig. 4B,E). In case of the D135N mutant, AS283 appeared to increase the ionomycin-induced movement (Fig. 4B,F), however, this was significant only by t-test and not by ANOVA (Fig. 4B). Compared to WT, the movement of the D135N mutant in the presences of AS283 was no longer significantly different by ANOVA, but was still significantly lower by t-test (Fig. 4B). Overall, the data in Fig. 4 show that ionomycin-induced movement of CaMKII to GluN2B in HEK cells is impaired by the D135N mutation, similar as shown for light induced movement of paCaMKII in Fig. 3. They also suggest that ionomycin-induced movement of the D135N mutant might be partially increased by AS283, but this is less clear than in case of the light induced movement of paCaMKII in Fig. 3. Additionally, in contrast to the light induced movement of paCaMKII, no impairment of ionomycin-induced movement was seen here for the T286A mutant, suggesting that ionomycin provides a stronger stimulus that creates a ceiling effect with maximal movement for both WT and T286A but not D135N mutant, which moved less than the T286A mutant in both assays.

**Figure 4:**
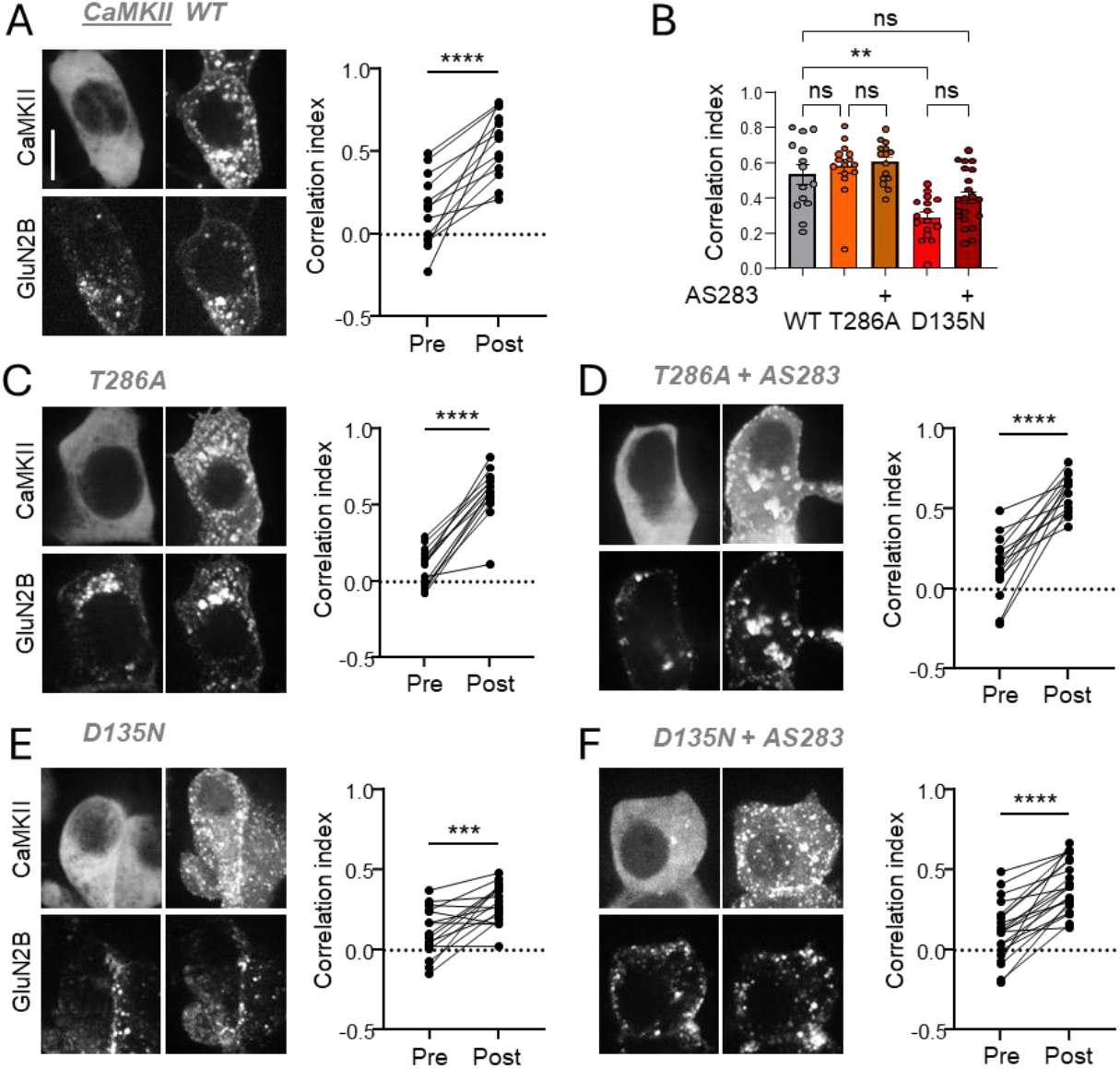
The D135N mutation reduces Ca^2+^-induced CaMKII movement to GluN2B in HEK cells. (**A**) Representative images of WT YFP-CaMKII and mCh-GluN2B pre- and post-ionomycin stimulation and quantification (n=13 cells, paired t-test). (**B**) Quantification of CaMKII-GluN2B correlation post-stimulation for WT, T286A, and D135N CaMKII with and without AS283 (n=14,15,15,16,21 cells, two-way ANOVA with Šídák’s multiple comparison’s test). (**C**) Representative images of T268A GFP-CaMKII and mCh-GluN2B pre- and post-ionomycin stimulation and quantification (n=15 cells, paired t-test). (**D**) Representative images of T268A GFP-CaMKII and mCh-GluN2B pre- and post-ionomycin stimulation following AS283 treatment and quantification (n=15 cells, paired t-test). (**E**) Representative images of D135N YFP-CaMKII and mCh-GluN2B pre- and post-ionomycin stimulation and quantification (n=16 cells, paired t-test). (**F**) Representative images of D135N YFP-CaMKII and mCh-GluN2B pre- and post-ionomycin stimulation following AS283 treatment and quantification (n=21 cells, paired t-test).

### The D135N mutation allows only transient co-condensation with GluN2B

Stimulation by Ca^2+^/CaM induces both binding and co-condensation of CaMKII and GluN2B (7, 19, 20). Thus, we decided to also test the effect of the D135N mutation in our cellular assay of co-condensation (Fig. 5). Similar to our binding assays, the co-condensation assay involves co-expression of fluorescently labeled CaMKII and GluN2B C-tail in HEK cells, however, in this case the GluN2B C-tail is not membrane targeted but soluble. Then, both GFP-CaMKII and the soluble mScarlet-GluN2B C-tail are dispersed throughout the cytoplasm, but form clusters after a Ca^2+^-stimulus is induced by ionomycin treatment (Fig. 5A). These co-condensates were maintained even after chelation of Ca^2+^ with EGTA (Fig. 5A), consistent with previous results that showed that maintenance of the co-condensates within cells is mediated by pT286 (7). The GFP-CaMKII D135N mutant also formed co-condensates with GluN2B, but these condensates were not maintained beyond removal of Ca^2+^ with EGTA (Fig. 5B), as expected based on the deficiency of this mutant to undergo pT286 (see Fig. 1). Direct comparison of the co-condensation of GFP-CaMKII wildtype versus D135N mutant in a time-lapse analysis illustrates the differential persistence during the EGTA treatment (Fig. 5C). Additionally, direct comparison shows that while D135N significantly co-condensates with GluN2B, the extent of co-condensation is significantly lower compared to CaMKII wildtype (Fig. 5C), similar as was seen in our other experiments for the binding.

**Figure 5:**
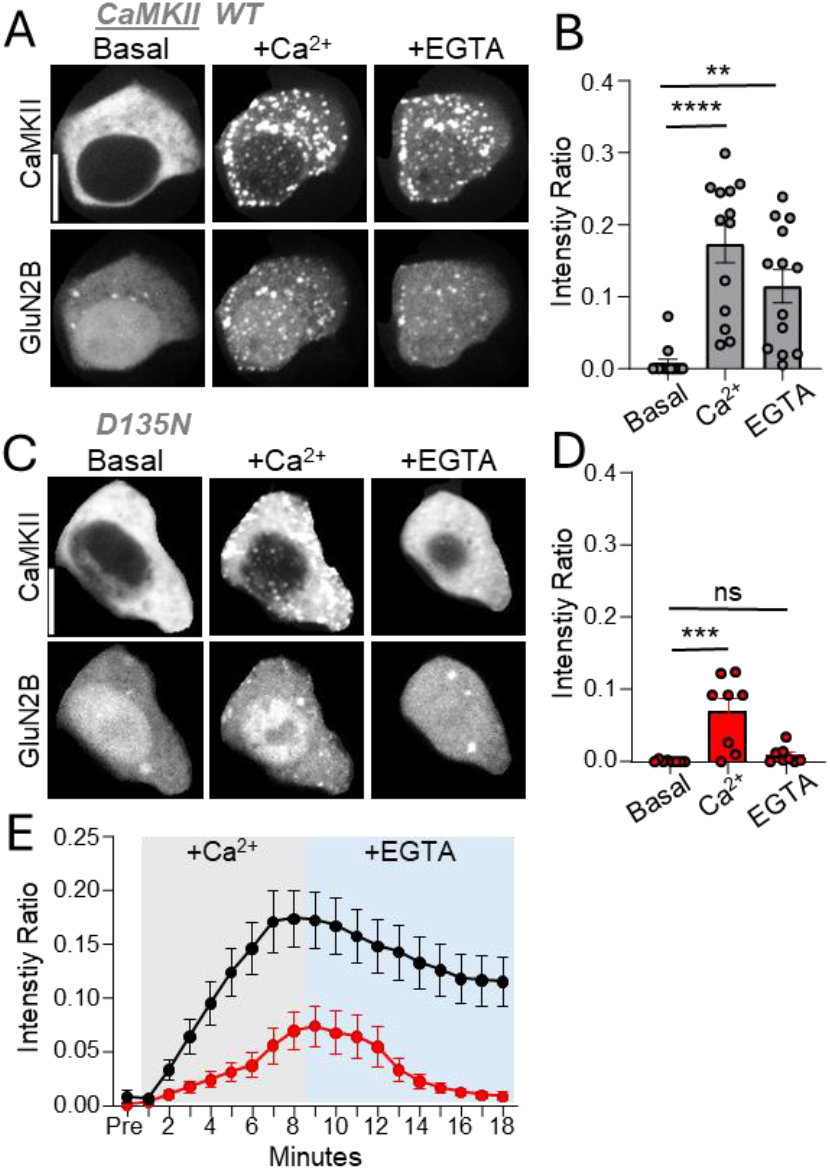
The D135N mutation reduced co-condensation of CaMKII with GluN2B and prevents its persistence beyond the Ca^2+^-stimulus. (**A**) Representative images of CaMKII WT and GluN2B under basal conditions (Basal), after Ca^2+^ stimulation with ionomycin (+Ca^2+^), and after Ca^2+^ chelation with EGTA (+EGTA). (**B**) Quantification of CaMKII WT/GluN2B co-condensation across the indicated conditions (one-way ANOVA with Tukey’s post-hoc test; **p<0.01, ***p<0.001). (**C**) Representative images of CaMKII D135N and GluN2B under the same conditions as in A. (**D**) Quantification of CaMKII D135N/GluN2B co-condensation (one-way ANOVA with Tukey’s post-hoc test; ns>0.5, *p<0.5) (**E**) Time course of CaMKII WT (black) and D135N (red) co-condensation before (Pre), during Ca^2+^ stimulation with ionomycin (gray), and after Ca^2+^ chelation with EGTA (blue). Co-condensation of WT CaMKII was induced by Ca^2+^ and maintained after EGTA, whereas co-condensation of D135N was fully reversed after EGTA. Scale bar, 10 μm.

## DISCUSSION

Here, we characterized the D135N tool mutant of CaMKII and found no detectable residual enzymatic activity. However, we found that the D135N mutation also caused reduced CaMKII binding to the cytoplasmic C-tail of GluN2B. Part of the observed reduction in binding within cells was expected due to the lack of enzymatic activity that mediates autophosphorylation at T286 (pT286), thereby abolishing one mechanism for positive regulation of CaMKII binding to GluN2B (15–17) and of the CaMKII movement to excitatory synapses in response to LTP stimuli that is mediated by the GluN2B binding (5, 7, 21). However, the reduction of GluN2B binding was more extensive for the D135N mutant than for a T286A mutant that also prevents pT286. Additionally, binding also appeared to be reduced in biochemical assays under conditions that prevented pT286 also in CaMKII wild type. This indicates that D135 may also directly contribute to the CaMKII interaction with GluN2B. Nonetheless, even though the D135N mutation significantly reduced GluN2B binding, it did not abolish the binding. Thus, the results with the D135N tool mutant are still consistent with the conclusion that GluN2B binding but not enzymatic activity of CaMKII is required for LTP induction (6). Notably, while consistent with this conclusion, the experiments with the D135N tool mutant tested the effects on basal transmission rather than on LTP (6). Nonetheless, the structural role in LTP induction was previously supported by several independent lines of evidence (5), with further evidence corroborating that the structural CaMKII function is indeed GluN2B binding (7). Thus, all three studies agree regarding the conclusion that enzymatic CaMKII activity is not required for LTP induction (5–7). However, one of these studies shows that enzymatic CaMKII activity is instead required for the subsequent phase of LTP expression (7), and that the required enzymatic activity is specifically the autonomous activity that is generated by CaMKII binding to GluN2B (7, 15). A more important discrepancy between the studies may be regarding LTP maintenance: One of the studies claims to show that pT286 is required for LTP maintenance (6), even though the LTP experiments shown do not distinguish between the different phases of LTP, which was also the case for the original study that first demonstrated pT286 requirement in LTP (22). Additionally, both other studies directly addressed the role of pT286 experimentally: They showed that the initial requirement for pT286 in LTP induction can be circumvented by ATP-competitive CaMKII inhibitors that directly enhance CaMKII binding to GluN2B (5, 7). This allowed LTP induction even in T286A mice. Most importantly, after washout of the drug, LTP was then expressed and maintained even in the absence of pT286 (5, 7). Thus, these results strongly indicate that pT286 is not required for LTP maintenance (4), consistent also with prior studies that indicated that pT286 at synapses last only for minutes after LTP stimuli (23). Several other studies have suggested that LTP maintenance may instead rely on CaMKII binding to GluN2B (24–26), however, such a role of GluN2B binding is still less well-supported than the more recently described roles in LTP induction and expression (4, 5, 7).

All of the three recent studies point to a central role of CaMKII binding to GluN2B in LTP (5–7) and this is consistent also with multiple other studies that were published over the last 15 years (24–27). However, this notion appeared to require reconciliation with three classic studies that showed that constitutively active monomeric forms of CaMKII are sufficient to induce synaptic potentiation (28–30) even though these monomers are impaired for binding to GluN2B (17, 31, 32). One possibility was that residual GluN2B binding of the monomers is sufficient to mediate potentiation even though it was not sufficient to mediate detectable movement to GluN2B within cells (31, 32). Another possibility was that the monomeric CaMKII induces synaptic potentiation by mechanisms that are different from LTP, even though they occlude subsequent further potentiation by LTP stimuli (28, 29). Our recent results instead suggest that monomeric CaMKII can circumvent requirement for GluN2B binding if it is artificially made enzymatically active in a Ca^2+^-independent manner that is also phosphatase resistant (33), i.e. is made autonomous in a manner that is normally provided by GluN2B binding by not by pT286. Clarification of these different possibilities further supports the functions of enzymatic activity of GluN2B-bound CaMKII in LTP, but this had to be achieved by tools other than the D135N mutant that was characterized in the current study.

## EXPERIMENTAL PROCEDURES

### Material availability

Requests for resources, reagents, or questions about methods should be directed to K. Ulrich Bayer. This study did not generate new unique reagents.

### Material and DNA constructs

Material was obtained from Sigma, unless noted otherwise. Expression of the intrabody is driven by the CAG promoter; expression of all other constructs is driven by the CMV promoter. The construct for expression of mCherry-CaMKII was made based on an mGFP-CaMKIIα construct (31) by switching the mGFP for an mCherry using the Eco47III and BsrgI sites. All constructs were validated by sequencing.

### Assays of CaMKII activity and autophosphorylation

CaMKII kinase activity was assessed by phosphate incorporation into peptide substrates (13, 31). Assays were done at 30oC for 1 min and contained HEK cell extracts diluted to contain equal amounts of total protein and 2.5 nM CaMKIIα kinase subunits (WT, K42M or D135N), 50 mM PIPES, pH 7.1, 0.1% BSA, 1 mM CaCl2, 1 µM Calmodulin, 10 mM MgCl2, 100 µM [γ-32P]ATP (~1mCi/mmole), and 75 µM syntide-2 substrate peptide. Kinase reactions were stopped by spotting onto P81 cation exchange chromatography paper (Whatman) squares and immersing in 0.5% phosphoric acid. After extensive washes in water, phosphorylation of the substrate peptide bound to the P81 paper was measured by liquid scintillation counting in Bio-Safe NA counting cocktail (RPI).

To determine phosphorylation of pT286, HEK cell extracts expressing WT, T286A or D135N CaMKIIα were diluted to contain equal amounts of total protein and 30 nM CaMKIIα. T286 was autophosphorylated in the above reaction buffer that contained 1 mM ATP and lacked [γ-32P]ATP and syntide. The reactions were carried out for 15 seconds, 1 min or 10 min at 30 ºC or on ice. Reactions were stopped by diluting into Laemmli buffer with 5 mM EDTA added, then boiled for 5 minutes. Phosphorylation of pT286 was determined by western blot.

### Western blots

Western blots were performed as described previously(5, 27). Before undergoing SDS-PAGE, the samples were heated in a sample buffer (final concentrations of 54 mM Tris pH 6.8, 1.6% SDS, 8% glycerol, 50 mM dithiothreitol, and 0.013% bromophenol blue) for 5 minutes at 95 °C. Proteins were separated using gradient precast gels (BioRad) and then transferred to low-fluorescence polyvinylidene difluoride membranes for 1 hour at 4 °C. Membranes were blocked in a fluorescent blocking buffer (Azure) and then incubated with anti-CaMKII (1:1000, BD Biosciences) or anti-CaMKIIα (1:2000, CBα2, made in house), anti-GST (1:1000, Millipore), and anti-pT286 CaMKII (1:3000, Phosphosolutions). After incubation with fluorescent secondary antibodies, antimouse AzureSpectra 700 (1:10,000, Azure Biosciences) or antirabbit AzureSpectra 800 (1:10,000, Azure Biosciences) signal was measured by fluorescent imaging using an Azure Western Blot Imaging System. Densitometric quantification was performed using Fiji ImageJ.

### In vitro GluN2B binding assay

CaMKII binding to the GluN2B C-terminal tail was performed in our established binding assay (7, 15). Anti-GST antibody–coated 96-well microtiter plates (Thermo Scientific) were washed three times in wash buffer (PST: 50 mM PIPES pH 7.12, 150 mM NaCl, 0.1% Tween-20). Wells were coated with saturating amounts of GST–GluN2B C-tail (GST–GluN2Bc, amino acids 1122– 1482) diluted in PST containing 0.05% BSA for 1 h at room temperature under gentle agitation, followed by three washes in PST. Plates were blocked with 5% BSA in PST for 30 min at room temperature. CaMKII (50 nM subunits; YFP2–CaMKII WT or D135N from HEK293T cell lysates) was quantified against a purified CaMKII standard in two rounds: first to determine the initial lysate concentration, and then after adjusting lysates to equal concentrations, which were re-verified against the standard to ensure accuracy. For binding, lysates were incubated for 30 min in kinase binding buffer (50 mM PIPES pH 7.12, 150 mM NaCl, 1 mM MgCl2, 0.1 mM ADP, 0.05% BSA, 0.05% Tween-20) containing Ca^2+^/CaM (2 mM/1 μM) or inhibitors as indicated. Wells were washed three times in PST and bound proteins were eluted in gel loading buffer (1.6% SDS) for 10 min at 95 °C, bound CaMKII was measured via immunoblot.

### Cell culture of HEK293 cells

Human embryonic kidney cells (HEK293; authenticated by short tandem repeat analysis) were cultured in Dulbecco’s modified Eagle’s medium (Gibco) supplemented with 10% foetal bovine serum (Sigma) and 1% penicillin–streptomycin solution (Gibco). HEK cells were not tested for mycoplasma. HEK cells were grown on 10 cm culture flasks and split every 3–4 days (at approximately 90% confluency). For imaging experiments, cells were split into 12-well culture dishes on 18 mm no. 1 glass coverslips.

### Live imaging of HEK cells

HEK cells were grown and transfected with expression vectors for GFP or YFP–CaMKII mutants and pDisplay-mCh-GluN2B-c tail (2Bc) as previously described; the GluN2B construct used included an S1303A mutation to avoid potential complications by differential phosphorylation of this regulatory site in the different conditions with or without kinase inhibitors. GFP/YFP–CaMKII colocalization with GluN2B in response to a Ca^2+^ stimulus induced by 10 μM ionomycin was monitored for 5–10 min at 32 °C in imaging buffer (0.87× Hanks Balanced Salt Solution, 25 mM HEPES, pH 7.4, 2 mM glucose, 2 mM CaCl_2_, 1 mM MgCl_2_) by fluorescence microscopy. Images were acquired on a Zeiss Axiovert 200M equipped with a climate control chamber using SlideBook software (Intelligent Imaging Innovations; v.2025.0). Colocalization analysis was performed by calculating the Pearson’s correlation (correlation index) with ImageJ of pDisplay-mCh-2Bc and GFP–CaMKII within the cytoplasm of HEK cells after background subtraction.

### paCaMKII stimulation in HEK cells

HEK cells were transfected with GFP–paCaMKII (WT, T286A and D135N) and pDisplay-mCherry-2BC^S1303A^ and left in the dark for 10 min before photoactivation to ensure that paCaMKII was in the dark state. GFP–paCaMKII was then photoactivated and imaged simultaneously via confocal imaging over a 3 μm Z stack (step size: 0.6 μm) or a single plane with 488 nm excitation once per minute for a total of 5 min. The correlation index was measured the same as for ionomycin-induced colocalization.

### Assay of CaMKII co-condensation with GluN2B in HEK293T cells

HEK cells were transfected and imaged as described above. However, in this case, mEGFP-CaMKII was co-expressed with a soluble, nonmembrane-targeted version of the mScarlet-GluN2B-c tail. Co-condensate formation was induced by triggering a Ca^2+^ stimulus with 10 μM ionomycin for 6 minutes; dispersal or maintenance of the co-condensates was monitored after subsequent chelation of Ca^2+^ with 2.5 mM EGTA for an additional 10 minutes at 34 °C in an imaging buffer. For analysis, the CaMKII intensity ratio, defined as the fluorescent intensity within CaMKII puncta divided by the total fluorescent intensity within the cell per time point, was used to quantify the dynamics of co-condensate formation and maintenance. As the level of co-clustering was negatively correlated with the level of CaMKII expression, we eliminated low and high-expressing cells from the analysis in order to ensure that the 2 conditions were compared in cells with similar medium levels of CaMKII expression.

### Quantification and statistical analysis

All data are shown as mean ± SEM. Statistical significance is indicated in the figure legends. Statistics were performed using Prism (GraphPad) software. Imaging experiments were obtained and analyzed using SlideBook 2025 software. All comparisons between two groups met parametric criteria, and independent samples were analyzed using unpaired, two-tailed Student’s t-tests. Comparisons between three or more groups meeting parametric criteria were done by one-way ANOVA with specific post-hoc analysis indicated in figure legends. Comparisons between three or more groups with two independent variables were assessed by two-way ANOVA with Bonferroni post-hoc test to determine whether there is an interaction and/or main effect between the variables. Asterisks represent level of significance: *p<0.05;**p<0.01; ***p<0.001.

## DATA AND CODE AVAILABILITY

The datasets generated during this study are available through Mendeley. No original code was generated during this study.

## AUTHOR CONTRIBUTIONS

M.E.L., C.M.B., R.V. and S.J.C. performed experiments and analyses; M.E.L., C.M.B. and K.U.B. conceived this study; K.U.B. wrote the initial draft and all authors contributed to the final manuscript.

## FUNDING AND ADDITIONAL INFORMATION

This work was supported by National Institutes of Health grants F31 AG084197 (to M.E.L.), R01 NS081248, R01 NS118786, and R01 AG067713 (to K.U.B.).

